# Cas9HF1 enhanced specificity in *Ustilago maydis*

**DOI:** 10.1101/671826

**Authors:** Weiliang Zuo, Jasper RL Depotter, Gunther Doehlemann

## Abstract

The clustered regularly interspaced short palindromic repeats (CRISPR)-Cas9 system is widely used as a tool to precisely manipulate genomic sequence targeted by sgRNA (single guide RNA) and is adapted in different species for genome editing. One of the major concerns of CRISPR-Cas9 is the possibility of off-target effects, which can be remedied by the deployment of high fidelity Cas9 variants. *Ustilago maydis* is a maize fungal pathogen, which has served as a model organism for biotrophic pathogens for decades. The successful adaption of CRISPR-Cas9 in *U. maydis* greatly facilitated effector biology studies. Here, we constructed an *U. maydis* reporter strain that allows *in vivo* quantification of efficiency and target specificity of three high fidelity Cas9 variants, Cas9HF1, Cas9esp1.1 and Cas9hypa. This approach identified Cas9HF1 as most specific Cas9 variant in *U. maydis*. Furthermore, whole genome sequencing showed absence of off-target effects in *U. maydis* by CRISPR-Cas9 editing.

## Introduction

The CRISPR-Cas9 is part of the bacterial immune system to fend off bacteriophage infection, which was first identified in *Streptococcus pyogenes* (Barrangou et al. 2007). The Cas9 protein serves as endonuclease and induces double strand breaks in the targeted region (Jinek et al. 2012) where the specificity of Cas9 is determined by the loaded guide RNA (gRNA) containing the first 20 nt spacer which is complement to the sequence of gene-of-interest with a protospacer adjacent motif (PAM) (Gasiunas et al. 2012). The double strand break is then repaired by the error-prone non-homologous end joining (NHEJ) or homologous direct repair. The capacity of CRISPR-Cas9 to manipulate defined genomic targets in addition to the easy design and manipulation makes it a powerful gene editing tool to create gene knockouts or conversion (Charpentier and Doudna 2013; Jiang and Doudna 2017). Currently, CRISPR-Cas9 based technologies are also widely used in various applications such as activation of gene expression, genomic labeling (Chen et al. 2013, 2016; Ma et al. 2015; Tanenbaum et al. 2014), epigenetic modification (Liao et al. 2017; Pulecio et al. 2017) and translational disruption (Pulecio et al. 2017). However, a major concern of CRISPR-Cas9 technology is the possibility of off-target effects, where sequences shared high similarity to the target are also cleaved by Cas9 (Fu et al. 2013; Hsu et al. 2013; Kosicki, Tomberg, and Bradley 2018).

To solve this problem, several strategies were applied, including using lower levels of active Cas9 (Davis et al. 2015; Pulecio et al. 2017), shortening the gRNA sequence at the 5’-end region (Fu et al. 2014), producing Cas9 nickase mutant or a Cas9 nuclease mutant fused with a FokI nuclease domain (Fu et al. 2014; Guilinger, Thompson, and Liu 2014). However, these methods often compromise the on-target efficiency or complicate the cloning process. Additionally, high fidelity Cas9 variants were generated such as Cas9HF1 (Slaymaker et al. 2016), Cas9esp1.0, 1.1 (Kleinstiver et al. 2016) and Cas9hypa (Chen et al. 2017), which demonstrate enhanced specificity without reducing on-target efficiency.

*Ustilago maydis* is a pathogenic fungus that causes smut disease in maize. It can infect all the above-ground tissues of maize plants and induces local tumor formation within two weeks after infection under glasshouse conditions (Kämper et al. 2006). Compared to other smut fungi, the unique and rapid tumor development makes it an excellent model to study biotrophic plant pathogens. The pathogenicity of *U. maydis* is initiated by the recognition and fusion of different mating strains, which is accompanied by morphological switch from yeast like growth of haploid sporidia to a diploid filament (Bölker, Urban, and Kahmann 1992; Spellig et al. 1994). The assembly of compatible mating genes in one single genetic background to create the solopathogenic strain SG200 facilitates pathogenic development of *U. maydis* without prior mating (Kämper et al. 2006).

Similar to other plant pathogens, the virulence of *U. maydis* is largely determined by its repertoire of secreted effector proteins, which are mostly highly expressed during host infection to trigger fungal growth and cause disease (Skibbe et al. 2010). Only few individual effector genes with large effect on virulence have been functional characterization (Djamei et al. 2011; Doehlemann et al. 2009; Ma et al. 2018; Mueller et al. 2013; Redkar et al. 2015; Tanaka et al. 2014). However, many effectors are present in gene families and / or show functional redundancy, which requires deletion of multiple genes at the same time to obtain visible virulence defects (Zuo et al. 2019, in press). An FLP (flippase)-recombinase based system for marker rescue allows multiple gene deletions in *U. maydis*, however is limited by the potential genome rearrangement between remaining FRT (flippase recognition target) sequences in the genome and time consuming process (Khrunyk et al. 2010). To make use of the significant advantages of CRISPR-Cas9 over classical homologous recombination, Schuster et. al adapted the CRISPR-Cas9 system in *U. maydis* by generating a codon optimized Cas9 protein and expression of the sgRNA under control of the *U. maydis* U6 promoter, which allowed high efficiency in genome editing (Schuster et al. 2016). Furthermore, tRNA promoters were used for multiplexing sgRNAs, empowering knockouts of multiple genes to be generated by one construct (Schuster, Schweizer, and Kahmann 2018). The CRISPR-Cas9 efficiency in *U. maydis* was further improved up to 40-100% by expressing Cas9 under *U. maydis* heat shock protein 70 promoter even with sgRNA has 20th PAM-proximal mismatch (Schuster et al. 2018). However, this brought concerns of how to increase Cas9 specificity in *U. maydis* without sacrifice the high efficiency.

In this study, we generated a *U. maydis* reporter strain expressing green fluorescent protein (GFP) in an expression cassette flanked by designed off-targets with 19th PAM-proximal mismatch sequence for a *bw2* sgRNA. Using this reporter strain, we found that Cas9HF1 confers significantly increased fidelity in *U. maydis* when compared to Cas9wt and other Cas9 variants. Furthermore, by Illumina-sequencing we detected no off-target effect in the *U. maydis* genome by testing two different sgRNAs by CRISPR-Cas9 editing with of the tested Cas9 versions.

## Results

### Construction of GFP reporter strain for off-target screening

In previous studies, two main strategies were used to test specificity of Cas9 variants. One is use sgRNAs have been reported to have off-target effects and monitor these known off-targets sites in the mutants to evaluate the specificity of Cas9 variants (Chen et al. 2017; Kleinstiver et al. 2016; Slaymaker et al. 2016), the other approach is to conduct gene knockouts by using sgRNAs with different mismatched nucleotides to the target and then detected the editing efficiency (Kim et al. 2017; Zhang et al. 2017). Although CRISPR-Cas9 was adapted in *U. maydis*, there are few publications on the application of this technology for gene deletion, not to mention the discovery of sgRNAs with off-target effect confirmed by whole genome sequencing or *in-vitro* test.

To test the high specificity Cas9 variants, we selected the *bw2* gene as target for genome editing as it has been done previously (Schuster et al. 2016), however, we designed one sgRNA in which the entire 20 nt spacer is matched to the target (including the first G required for the transcription under U6 promoter) **(Fig. 1a)**. In order to increase the throughput and facilitate the evaluation of on-target and off-target effect at the same time, a GFP reporter strain SG200-19MM was generated based on the solopathogenic *U. maydis* strain SG200. The GFP was expressed under control of the o*tef* promoter, which confers strong expression under axenic culture growth conditions. The expression cassette was flanked by two designed off-targets, which contained a single nucleotide mismatch at 19th PAM-proximal position compares to the designed *bw2* sgRNA **(Fig. 1a).** The cassette was integrated into the *ip* (iron-sulphur protein) locus (Broomfield and Hargreaves 1992) of SG200 by homologous recombination **(Fig. 1a)**. Single copy integration into the *ip* locus was confirmed by southern blot (not shown). The resulting strain SG200-19MM showed a stable GFP signal in induced filaments on charcoal PD plates (**Fig. 1b, c**). Our reporter screen is based on two readouts: on-target disruption of the *bw2* gene in SG200-19MM causes loss of filamentous growth on charcoal PD plates (fuzz- phenotype, in case of the frame-shift) (**Fig. 1b, c**). In addition, double strand break on the 19MM off-target will result in the loss of the GFP signal due to the cleavage of the GFP expression cassette from the genome as consequence of off-target editing **(Fig. 1b, c)**.

**Fig1.**
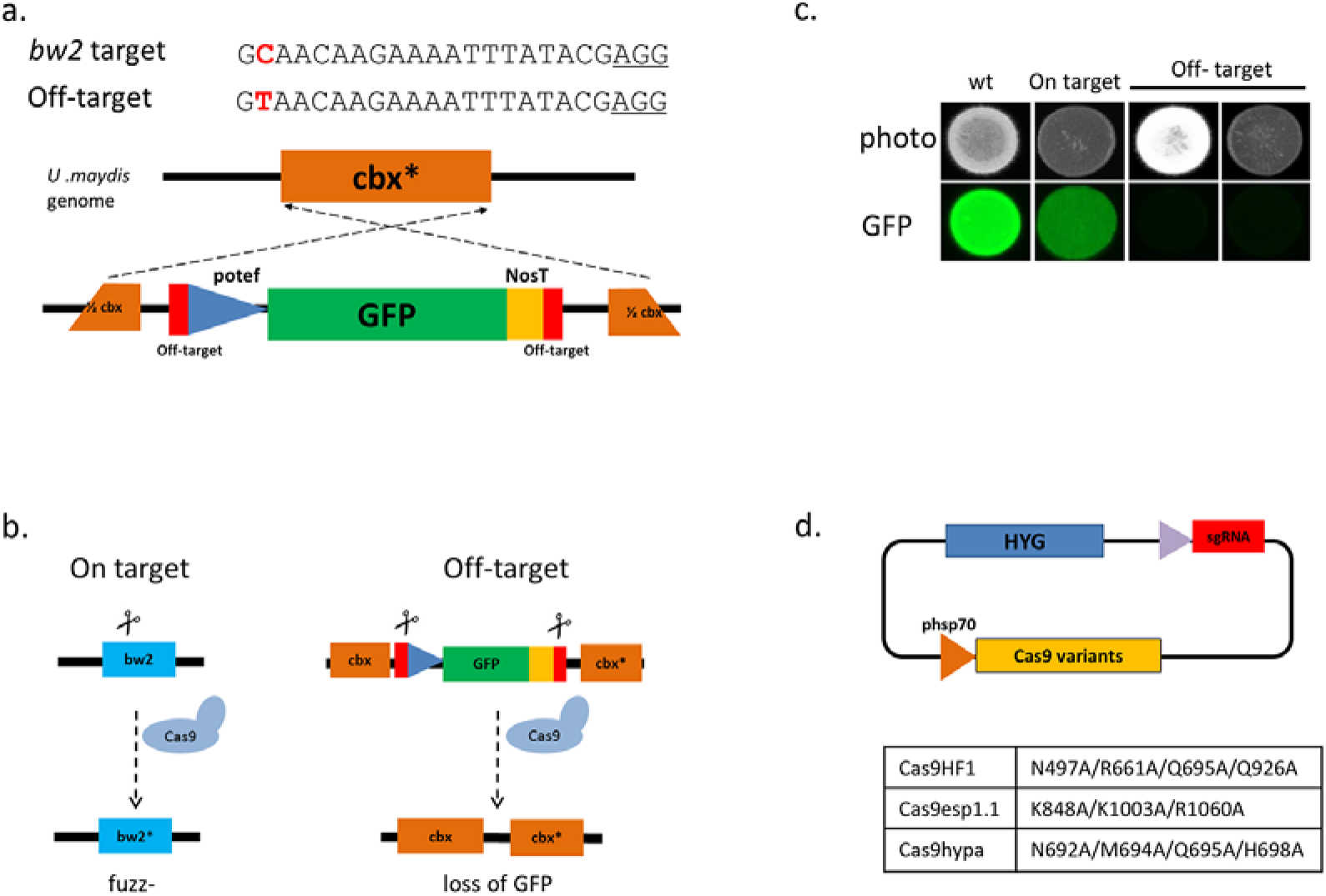
Construction of SG200 GFP reporter strain and test of Cas9 high specific variants. **a)** Scheme showing the construction of reporter strain SG200-19MM. *bw2* sgRNA and its 19th PAM-proximal off-targets sequences were shown. The red font indicated the nucleotide change between on and off targets, and the PAM sequences were underlined. The off-targets were inserted into the flank region of GFP expression cassette and the GFP cassette was integrated into the *U. maydis ip* (succinate dehydrogenase iron–sulfur protein subunit) locus and resulted in carboxin resistance of SG200-19MM. **b)** Scheme showing the phenotype of on-target and off-target effects in the reporter strain. cbx*: carboxin susceptible allele in *U. maydis* which containing one amino acid change compared to the functional allele that make *U. maydis* susceptible to carboxin. ½ cbx indicate the vector was linearized by cut cbx resistance allele into two halves for the homolog recombination. **c)** Phenotypes of on-target and off-target genome editing. The photo showed the fuzz / fuzz- growth of wild type and *bw2* knockouts on charcoal PD plate, the GFP image showed the off-target editing loss the GFP signal. **d)** The plasmids used for off-target testing and the corresponding mutations in the Cas9 variants tested. The Hygromycin resistance was used for selection on the free circulating plasmid containing Cas9.

### Specificity of Cas9 variants in *U. maydis*

Three high fidelity Cas9 variants, Cas9HF1, Cas9esp1.1 and Cas9hypa, were generated by inserting the required point mutations into the *U. maydis* codon optimized Cas9wt (Chen et al. 2017; Kleinstiver et al. 2016; Schuster et al. 2016; Slaymaker et al. 2016) (Fig1. c). The resulting Cas9 variants HF1, esp1.1 and hypa were then used to knock out the *bw2* gene in SG200-19MM **(Fig1. d)**. Transformants were first cultured in YEPS light medium overnight then dropped on charcoal PD plates to test the filament induction and detect the GFP signal. Four independent transformations were conducted, and each 46-48 independent colonies per treatment (23-24 for the first replicate) were tested on charcoal PD plate for phenotyping. The transformants that lost the GFP signal were considered to contain the off-target editing due to the cleavage of GFP expression cassette **(Fig. 1c)**. The off-target ratio was calculated by the number of colonies without GFP signal divided by the total number of colonies tested and compared between Cas9wt and the different the high fidelity variants. In all 4 independent experiments, Cas9HF1 resulted in consistently and significantly reduced off targeting by 8.97-25.72% compared to Cas9wt **(Fig. 2a, b)**. We next compared the fuzz-rate of transformants, which reflects the successful disruption of target *bw2* genes. Here, Cas9HF1 did not show any obvious compromised on-target efficiency compared to Cas9wt **(Fig. 2c)**. To our surprise, the other two Cas9 variants, Cas9esp1.1 and Cas9hypa did not enhance fidelity, but exhibited higher off-target effect compared to Cas9wt **(Fig. 2a, b)**, however the on target editing efficiency is not affected **(Fig. 2c, d)**. Based on this result, we identified Cas9HF1 as the most specific Cas9 variant without detectable reduction in on-target efficiency.

**Fig2.**
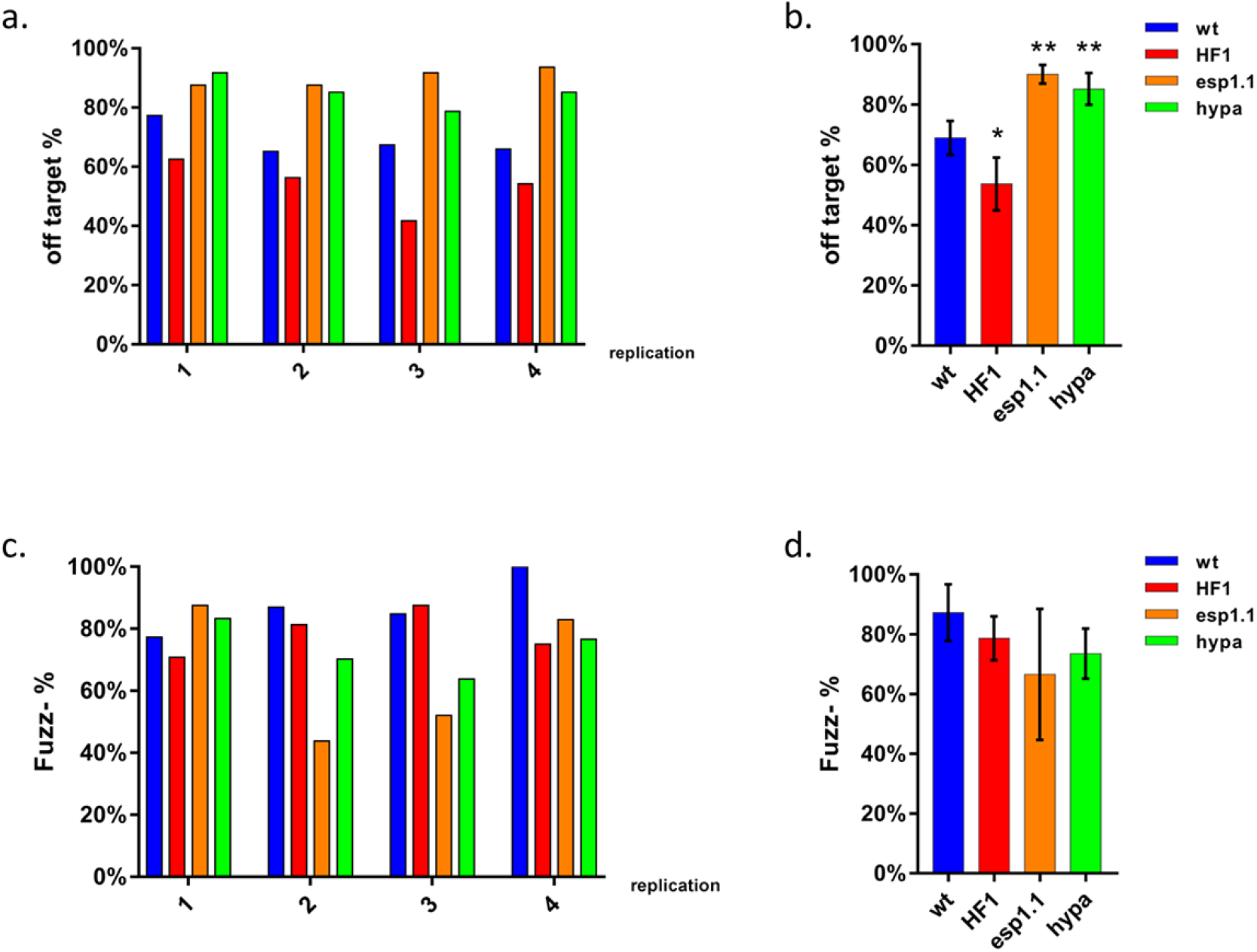
Evaluation of on-target and off-target efficiency of different Cas9 high specificity variants in the SG200-19MM reporter strain. **a)** The off-target rate of different Cas9 variants in *U. maydis*. Off-targeting was detected by the number of colonies lost GFP signal, and **b)** The summary result of 4 replicates, Cas9HF1 showed 8.97-25.72% significantly reduced off-targeting compared to wt, whereas the Cas9esp1.1 and Cas9hypa showed significantly higher off-targeting rate. **c)** On-target efficiency of Ca9 variants. The on target editing was revealed by the fuzz- colonies, which do not grow filamentous on charcoal PD plates. **d)** The summary of on target editing efficiency of different Cas9 variants. all three high fidelity Cas9 variant showed similar editing efficiency. Student *t-test* was used for statistical analysis. *, *p*<0.05. **, *p*<0.01.

### CRISPR-Cas9 off-target effects in *U. maydis*

In a next experiment, we performed whole genome re-sequencing to investigate whether CRISPR-Cas9 mediated gene knock-out causes any unexpected mutations during genome editing. In addition to the *bw2* gene, we targeted the *U. maydis fly1* gene which encodes a secreted fungalysin metalloprotease. Deletion of *fly1* results in reduced virulence and altered cell-separation of *U. maydis* in axenic culture (Ökmen et al. 2018). The CRISPR-Cas9 constructs expressing Cas9wt/Cas9HF1 together with sgRNAs for *bw2* or *fly1* were applied to strain SG200 separately. Genomic DNA of 8 independent colonies from each treatment were randomly chosen and subjected for Illumina sequencing. All transformants were tested to confirm the loss of CRISPR-Cas9 plasmid before DNA preparation. Before sequencing, the transformants were confirmed for successful on target editing by test of filament growth on charcoal PD plates for *bw2* knockouts, or T7 endonuclease I digestion assay for *fly1* knockouts, respectively. As a control, untransformed cells of the progenitor strain SG200 were sequenced to generate high quality reference genome.

In total, Illumina sequencing yielded between 8.4 and 12.6 million reads for the different samples resulting in an average gene coverage ranging between 45 and 66x. The SG200 reads were first mapped to the public available *U. maydis* reference genome “*U.maydis* 521” (Kämper et al. 2006) to create the SG200 reference, which excludes variations related to natural diversity between 521 and the SG200 strain used in our laboratory. The reads from the CRISPR-Cas9 transformants were then mapped to SG200 reference for variation calling. In total, we detected 78 deletions, 72 insertions and 225 SNVs (single nucleotide variant) from the CRISPR-Cas9 editing mutants **(Fig. 3a)**. Since the error prone NHEJ was considered to generate Indel in the genome, we excluded the SNVs from off-target analysis. From all the Indels identified, we also excluded the INDELs which were present in all mutants **(Fig. 3b)**. These all-present Indels have the same mutated sequence from all 4 different treatments compared to SG200 which implies these INDELs were spontaneous mutations in the SG200 cultures during protoplast preparation. In order to investigate whether the Indels were caused by off-target effect of CRISPR-Cas9 genome editing process, we used Cas-OFFinder (Bae, Park, and Kim 2014) to predict the possible off-targets of *bw2* and *fly1* sgRNAs in the *U. maydis* genome. A relaxed condition of “10 mismatches with one DNA/RNA bulge” was used as standard for prediction and none of these INDELs can be determined as off-target. Hence, we concluded that no off-target were generated during CRISPR-Cas9 genome editing.

**Fig3.**
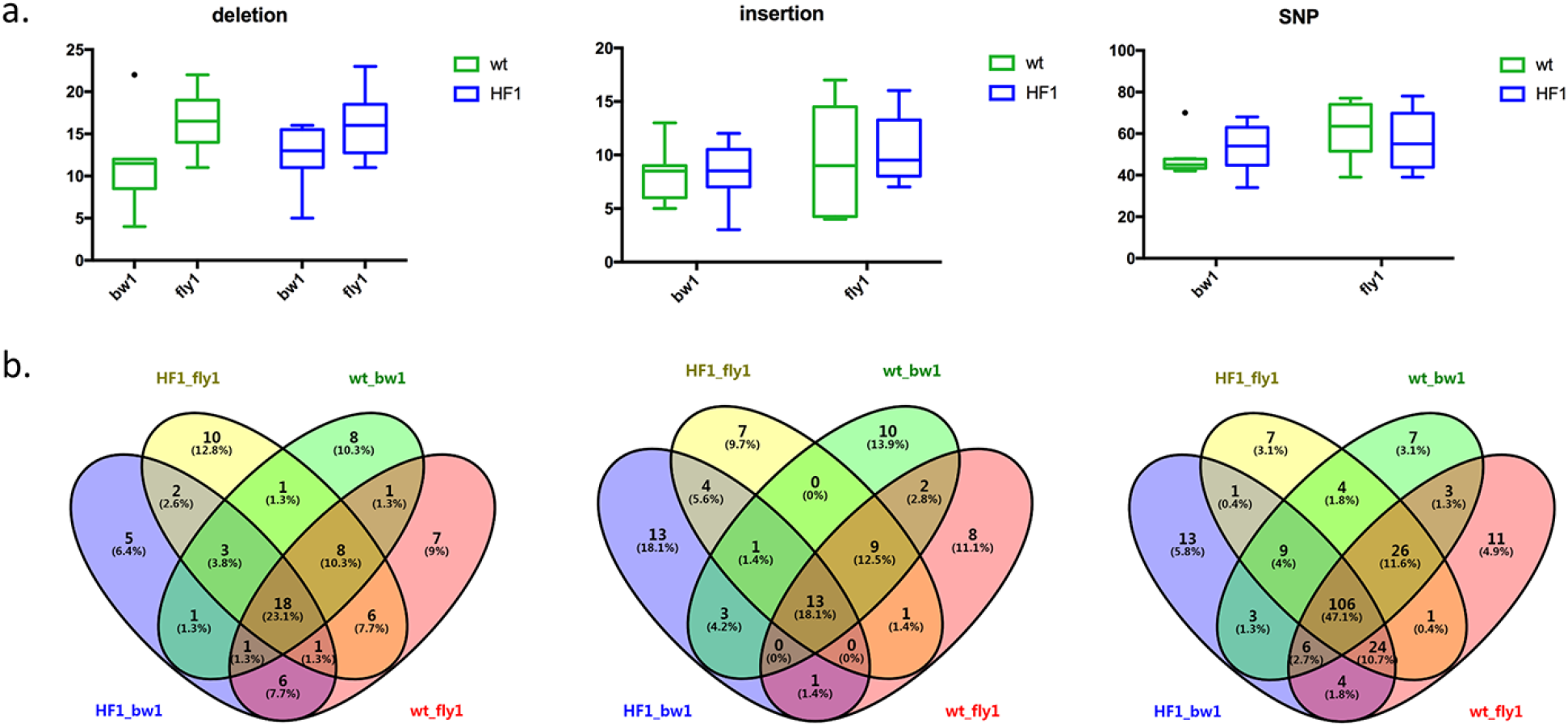
Whole genome sequencing of Cas9wt and Cas9HF1 editing mutants. **a)** Box-and-Whisker plot showed the total number of deletion, insertion and SNP identified from *bw2* and *fly1* knockouts. **b)** Venn diagram showed the distribution of mutations from different sgRNA and Cas9. 23.1% of deletions, 18.1% of insertions and 47.1% SNPs were detected from different treatment indicated they were generated during protoplast preparation.

## Discussion

To evaluate Cas9 specificity we generated a reporter strain SG200-19MM for fast *in vivo* detection of on target and off target editings simultaneously. The off-targeting in SG200-19MM simply determined by the loss of GFP signal due to the cleavage of GFP expression cassette in the genome which is flanked by two designed off-targets based on the *bw2* sgRNA. Previously, reporter strain systems facilitated off-target evaluation and helped to identified new Cas9 high specific variants in baker’s yeast (Casini et al. 2018). This *U. maydi*s SG200-19MM reporter strain was applied successfully in our study and can be used for testing any new emerging high specific Cas9 in future. Moreover, without the knowledge of known sgRNA causing off target mutations, this method is advantageous compared to testing different mismatched sgRNA for off-targeting. The sgRNA in such engineered cells will preferably bind to the 100% matched target over the mismatched off-targets, which might be more close to the native situation when unspecific genome editing is happening. In addition, using one sgRNA to detect on/off targeting at the same time eliminates the putative effect from the potentially variable difference amongst different sgRNAs sequences and different transformation events.

Our results showed that Cas9HF1 has an enhanced specificity compared to Cas9wt in *U. maydis*. Cas9 requires a minimal perfect match of the spacer to the target in the “seed region” (the first 8-12 PAM proximal nucleotides of the guide region) for cleavage (Semenova et al. 2011; Zhang et al. 2015), and the position of mismatch in the spacer affects the potential of off-targeting (Chen et al. 2017). We tested the 19th PAM proximal mismatch, which is more tolerated by Cas9, explaining the general high off-target events detected and small difference observed between Cas9HF1 and wildtype in the reporter strain assay. To our surprise, Cas9esp1.1 and Cas9hypa did not reduce off-target frequency over Cas9wt, but instead showed a higher rate of off-targeting. All these Cas9 variants were generated by the 3D structure based engineering method to change the energy requirements of the Cas9-sgRNA complex or sgRNA-target binding. This however could be affected by the intracellular environment of different species, which might be a possible explanation for the high unspecific targeting observed for Cas9esp1.1 and Cas9hypa.

In this study we could not observe any off-target activity Cas9wt and Cas9HF1 after the editing of the genes *bw2* and *fly2*. This is consistent with previous study in *U. maydis* (Schuster et al. 2016), although in this study we tested different sgRNAs and more independent colonies. The *U. maydis* genome is small, compact and largely lacks repetitive sequences, and together these features likely contribute to a low risk of Cas9-mediated off-site effects. Furthermore, the CRISPR-Cas9 module is transiently expressed in an autonomous replication plasmid, which can be quickly cleaned up from cell, short the interacting time of Cas9-sgRNA complex and genome. However, when multiplexing different sgRNAs in one construct required an elongated incubation to increase the life-time of CRISPR-Cas9 in the cell. While this might increase the chance of the off-targeting, use of Cas9HF1 in such experiments will greatly increase the specificity of editing, prevent the risk of off-targets.

## Materials and methods

### Strains and growth condition

The plasmids were transformed in *Escherichia coli* Top10 strains, and cultured in dYT liquid medium or YT plate with corresponding antibiotic. The solopathogenic *U. maydis* strain SG200 (*a1 mfa2 b1 bW2*) and SG200-19MM were cultured in YEPS light liquid medium or potato dextrose (PD, Difco) plate, or PD plate with 1% active charcoal for filament induction.

### Strain and plasmid construction

To generate strain SG200-19MM, the oligos containing the off-targets 5’-TATAGAACTCGAGCAGCTGA**GTAACAAGAAAATTTATACG<u>AGG</u>**AAGCTTGCATGCCTGCAGGTCG-3’ and 5’-CATGAGAATTCATCGATGAT**GTAACAAGAAAATTTATACG<u>AGG</u>**GATATCAGATCTGCCGGTCTCCC-3’ (off-target sequences were in bold, PAM sequences were underlined) were introduced into the flank region of GFP expression cassette in p123 plasmid in HindIII and EcoRV site sequentially by Gibson assembly (New England Biolabs, Ipswich, USA). The resulting plasmid p123-19MM was then linearized by SspI and transformed into SG200 protoplast as described previously (Schulz et al. 1990). The DNA of transformants was isolated and in-locus integration and copy number of insertions was confirmed by Southern blotting.

The Cas9 high fidelity variants were generated by “QuikChange Multi Site-Directed Mutagenesis Kit” (Agilent Technologies, CA, USA) with primers listed in **supplemental table 1**. To change the antibiotic resistance gene for the selection in *U. maydis*, the plasmids were digested with BsrGI, and integrated with hygromycin resistance cassette amplified from plasmid pUMa1507 (Terfrüchte et al. 2014) by Gibson Assembly (New England Biolabs, Ipswich, USA).

To construct the CRISPR vectors for gene knockout in *U. maydis*, sgRNAs were designed by E-CRISPR (http://www.e-crisp.org/E-CRISP/aboutpage.html) (Heigwer, Kerr, and Boutros 2014) (**supplemental table 1**). A similar approach was used for plasmid construction as described by Schuster et al. with some modifications (Schuster et al. 2016). In brief, 59 nt spacer oligomers containing the 20 nt “spacer” and 19 nucleotides (5’ upstream, overlap to plasmid) and 20 nucleotides (3’ downstream, overlap to scaffold) were ordered (Sigma, Darmstadt, Germany). The different Cas9 vectors were linearized with restriction enzyme Acc65I, and assembled with spacer oligo and “scaffold RNA” fragment with 3’ downstream 20 bp overlap to the plasmid by Gibson Assembly.

### Phenotyping and T7 endonuclease I digestion assay

The *bw2* gene knockout vectors were transformed into protoplasts of *U. maydis* strain SG200 or reporter strains SG200-19MM as previously described (Fotheringham and Holloman 1990). The transformants were transferred onto a new PD plate to grow overnight at 28°C, then the fresh colonies were picked and cultured in 300 μl YEPS light medium in 96-deep well plate with 200 rpm shaking at 28°C for 16-20 hours. 10 μl of overnight culture was dropped on charcoal PD plates for filament induction I and /or detection of GFP signal by ChemiDoc™ MP Imaging System (Bio-Rad, CA, USA). For the T7 endonuclease I assay, the genomic DNA of *fly1* knockouts and wildtype SG200 were prepared and a ~630 bp region containing the editing site was amplified by Phusion DNA ploymerase (New England Biolabs, Ipswich, USA) using the primers listed in **supplemental table 1**. Equal amount of wildtype and mutant PCR products were mixed and annealing to produce hybrid and digestion with 0.5 U T7 endonuclease I for 15 min at 37°C and then detected on agarose gel.

### Whole genome sequencing and off-target analysis

The *bw2* and *fly1* gene knockout mutants by Cas9wt and Cas9HF1 were cultured in YEPS light liquid medium overnight at 28°C, 200rpm. The DNA was prepared and purified by “MasterPure™ Complete DNA and RNA Purification Kit Bulk Reagents” (Epicentre, Wisconsin, USA). The DNA libraries were constructed using the Nextera DNA Flex Library Prep Kit, and paired-end sequencing was performed on the HiSeq4000 platform producing 75 bp long reads at the Cologne Center for Genomics (Cologne, Germany). Reads were checked for their quality with FastQC (v.0.11.6) and then used for further analysis (Andrews and Babraham Bioinformatics 2010) https://www.bioinformatics.babraham.ac.uk/projects/fastqc).

To create a SG200 *U. maydis* reference strain for read mapping, the assembly of strain 521 was used to map the SG200 sequence reads and subsequently call the variant to create a consensus strain (Kämper et al. 2006). Read-mapping was performed with the Burrows-Wheeler Aligner (BWA-MEM, v.0.7.17) (Li 2013). The variants (SNP and INDEL) were called using GATK after duplicate removal (McKenna et al. 2010). A new consensus genome, where the variants were implemented, was created using bcftools consensus (Narasimhan et al. 2016). This process of read-mapping and variant calling was iterated 9 times, so that a consensus strain was obtained were no variants could be called based on the SG200 reads. This consensus strain, hereafter called SG200 genome assembly, was then used as a reference to call variants to sequenced *U. maydis* strains that underwent mutagenesis through the CRISPR-Cas system. Similar as in the creation of the SG200 genome assembly, reads were mapped and variants were called with BWA-MEM (v.0.7.17) and GATK, respectively (Li 2013; McKenna et al. 2010). Only variants were called in genome regions were SG200 reads had coverage between 20-100x with the SG200 genome assembly. In addition, with the GATK VariantFiltration option the following requirements were set for variant calling: SNP = “QD < 2.0 ∥ FS > 60.0 ∥ MQ < 40.0 ∥ MQRankSum < −12.5 ∥ ReadPosRankSum < −8.0” and INDEL = “QD < 2.0 ∥ FS > 200.0 ∥ ReadPosRankSum < −20.0”. To see if variants corresponded to likely CRISPR-Cas off-target locations, off-targets were predicted in the SG200 genome assembly using Cas-OFFinder (Bae et al. 2014).

## Data availability

The genome sequencing data from Cas9wt and Cas9HF1 mediated *bw2* and *fly1* knockouts have been deposit in NCBI under the accession number PRJNA545211.

## Acknowledgements

We acknowledge funding from the European Research Council under the European Union’s Horizon 2020 research and innovation program (consolidator grant conVIRgens, ID 771035), as well as funding by the Deutsche Forschungsgemeinschaft (DFG, German Research Foundation) under Germany’s Excellence Strategy – EXC-2048/1 – Project ID: 390686111.

## Supplementary information

**Supplementary Fig1.** F**ilament growth induction and T7 endonuclease I assay to confirm the on-target editing in *bw2* and *fly1* knockouts respectively. a)** 10 µl culture of *bw2* knockouts were drop on charcoal PD plate to detected the ability of filament. Two drops from each 8 independent mutants were tested, the middle red circle indicated the SG200 control. **b)** T7 endonuclease I digestion of PCR product from *fly1* genes. The arrow indicated the expected big digest product after T7 endonuclease I digestion.

**Supplemental table 1. Primers used in this study.**

