## Supplementary figures and images for "Cas9HF1 enhanced specificity in *Ustilago maydis*"

### Supplemental Figure 1

a.

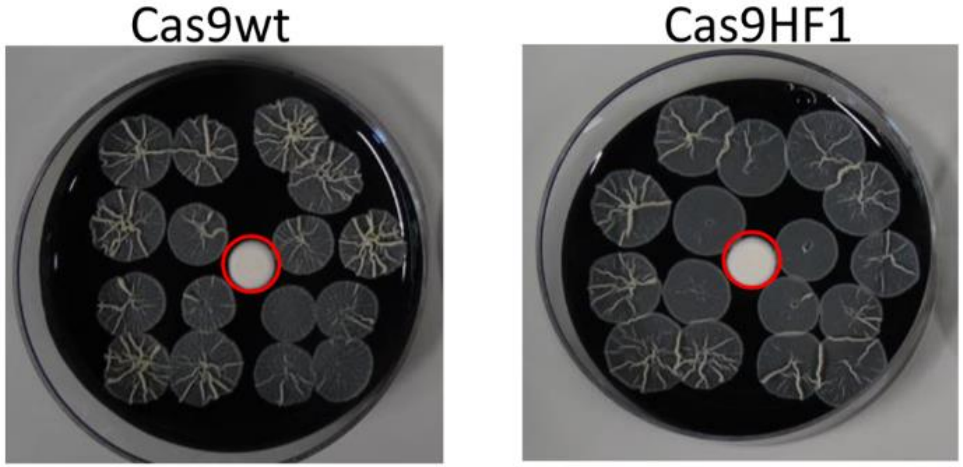


b.

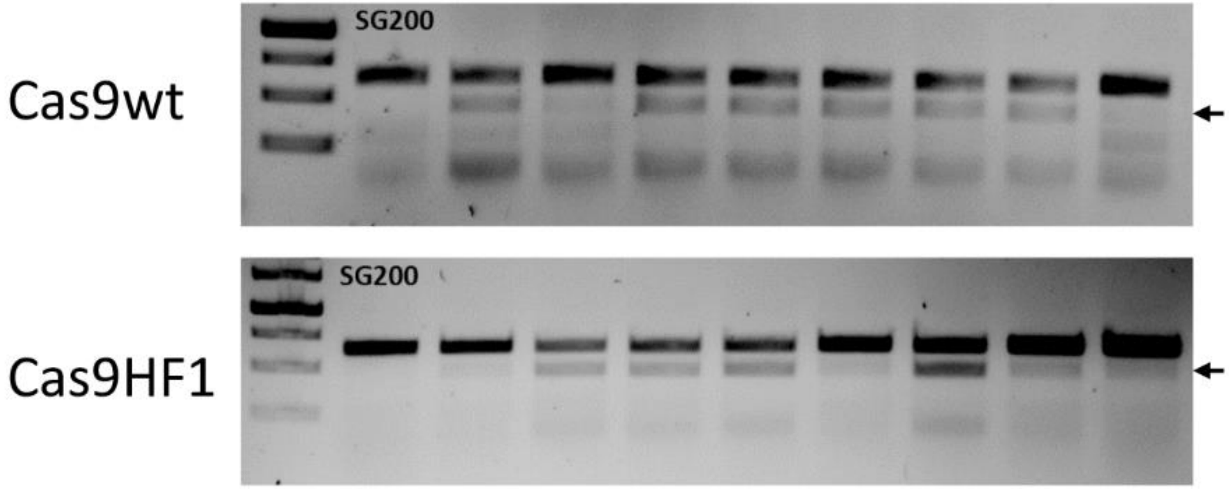
